# Pleiotropy promotes male exaggerated weapon and its associated fighting behaviour in a water strider

**DOI:** 10.1101/2020.01.09.898098

**Authors:** William Toubiana, David Armisén, Decaras Amélie, Abderrahman Khila

## Abstract

Exaggerated sexually selected traits, often carried by males, are characterized by the evolution of hyperallometry, resulting in their disproportionate growth relative to the rest of the body ^1–3^. While the evolution of allometry has attracted much attention for centuries, our understanding of the developmental genetic mechanisms underlying its emergence remains fragmented ^4,5^. Here we show that the hyperallometric legs in the males of the water strider *Microvelia longipes* are associated with a specific signature of gene expression during development. Using RNAi knockdown, we demonstrate that a broadly expressed growth factor, Bone Morphogenetic Protein 11 (BMP11, also known as Growth Differentiation Factor 11), regulates leg allometries through increasing the allometric coefficient and mean body size in males. In contrast, BMP11 RNAi reduced mean body size but did not affect slope in females. Furthermore, our data show that a tissue specific factor, Ultrabithorax (Ubx), increases intercept without affecting mean body size. This indicates a genetic correlation between mean body size and variation in allometric slope, but not intercept. Strikingly, males treated with BMP11 RNAi exhibited a severe reduction in fighting frequency compared to both controls and Ubx RNAi-treated males. Overall, we demonstrate a genetic correlation between male body size, the exaggerated weapon, and the intense fighting behaviour associated with it in *M. longipes*. Our results provide evidence for a role of pleiotropy in the evolution of allometric slope.

## Introduction

Extravagant ornaments and weapons, often carried by males in a wide range of lineages, represent some of the most striking outcomes of directional sexual selection driven by female choice or male competition ^2,3,6,7^. Exaggerated sexually selected traits are often characterized by their extreme growth and hypervariability among individuals of the same population ^8,9^. These features derive from changes in scaling relationships, especially the elevation of the allometric coefficient (i. e. hyperallometry), which results in certain structures growing disproportionately larger relative to the rest of the body ^8–11^. Despite the longstanding interest in the study of allometry, the genetic architecture underlying variation in allometric coefficient is still largely unknown ^9,12–15^. Allometric slope is known to be relatively stable during evolution and several studies suggest that this stasis could be the consequence of genetic correlation with other traits ^4,12–14,16,17^. In the context of trait exaggeration (e. g. hyperallometry), Fisher predicted a genetic correlation between male exaggerated ornaments and female preference for the exaggeration – a process known as runaway selection ^2^. Other studies, for example in the ruffs or horned beetles, also suggest a genetic link between the elaboration of male exaggerated traits and other secondary sexual traits such as male body size and reproductive behaviour ^18–20^. However, empirical tests of these predictions remain difficult to achieve. Determining the genetic architecture of allometric slope and testing its correlation with other traits would therefore greatly advance our understanding of the evolution of scaling relationships ^9,12–15^.

We address these questions in the water strider *Microvelia longipes*, where some males exhibit extremely long third legs due to a hyperallometric relationship with body size ^21^. Among the over 200 known species in this genus, *M. longipes* is the only species found where males simultaneously exhibit large body size, and high variation in both body and leg length ^21,22^. Males of other *Microvelia* species exhibit some but not all of these features, suggesting that the evolution of hyperallometry may be linked to the evolution of increased mean and variation in body size ^21^. In addition, males of *M. longipes* use the exaggerated legs to fight and dominate egg-laying sites where they intercept and mate with gravid females ^21^. Compared to its closest related species *M. pulchella*, *M. longipes* males fight over an order of magnitude more frequently, thus indicating that the evolution of the exaggerated weapon is also associated with the increase in competition intensity between males ^21^. Here we use comparative transcriptomics, along with studies of gene function and behaviour, to determine the genetic basis of third leg exaggeration in this species. We show that the developmental gene *Bone Morphogenetic Protein 11* (*BMP11*), also known as *Growth Differentiation Factor 11 (GDF11)*, plays a pleiotropic role both in the hyperallometry of the third leg, increase body size and fighting intensity in males, thus revealing a strict genetic correlation between these traits in the male of *M. longipes*.

## Results

### A burst of leg growth during the last nymphal instars

In *M. longipes*, third leg length is highly variable among males and this variability is higher in males compared to females (Figure 1A) ^21^. A developmental growth curve (Figure 1B; Supplementary figure 1) revealed that leg length is identical between individuals at the 1^st^ to 3^rd^ instars, but starts to diverge between the sexes and also among males at the 4^th^ instar (Figure 1B). A dramatic increase in leg length was observed between 5^th^ instar and adult males (Figure 1B). These differences in leg length between individuals are independent of any variation in developmental time from 1^st^ instar to adult (ANCOVA, F-value= 1.965, p-value= 0.1712; Supplementary figure 2). Therefore, the growth burst and its fluctuation at the end of post-embryonic development in male exaggerated legs accounts for both the sexual dimorphism and the hypervariability among males (Figure 1A-C). Ontogenetic allometry also confirmed the exaggerated growth rate of male third legs during development (Supplementary table 1). The first and second legs in males and females showed similar growth dynamics, although not as extreme as the third legs (Figure 1B). Based on these observations, we hypothesize that the striking variation in leg growth dynamics within and between the sexes is attributable to variation in gene expression at the end of nymphal development.

**Figure 1:**
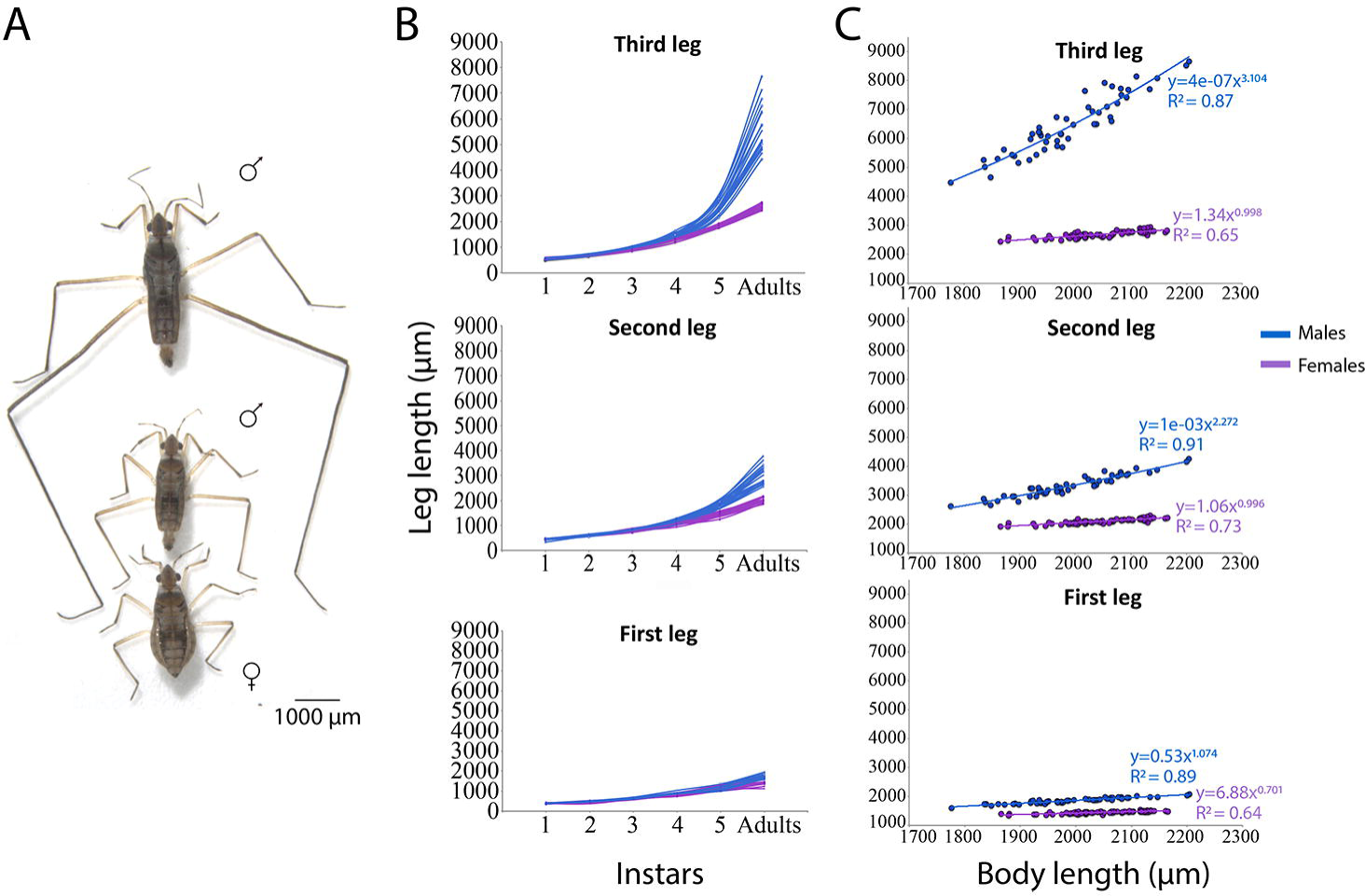
Growth dynamics of male exaggerated third legs in *M. longipes*. **(A)** Adult large and small males and a female showing final growth. **(B)** Leg length variation during post-embryonic development (nymphal development) in males and females. Each line represents leg growth dynamics of a single individual during nymphal development until adulthood. **(C)** Static allometries between body length and the three pairs of legs in males and females. Power low regressions were fitted to the raw data to represent the static allometric equation y=bx^a^. Differences in static allometry parameters were tested on log-transformed data in^21^.

**Figure 2:**
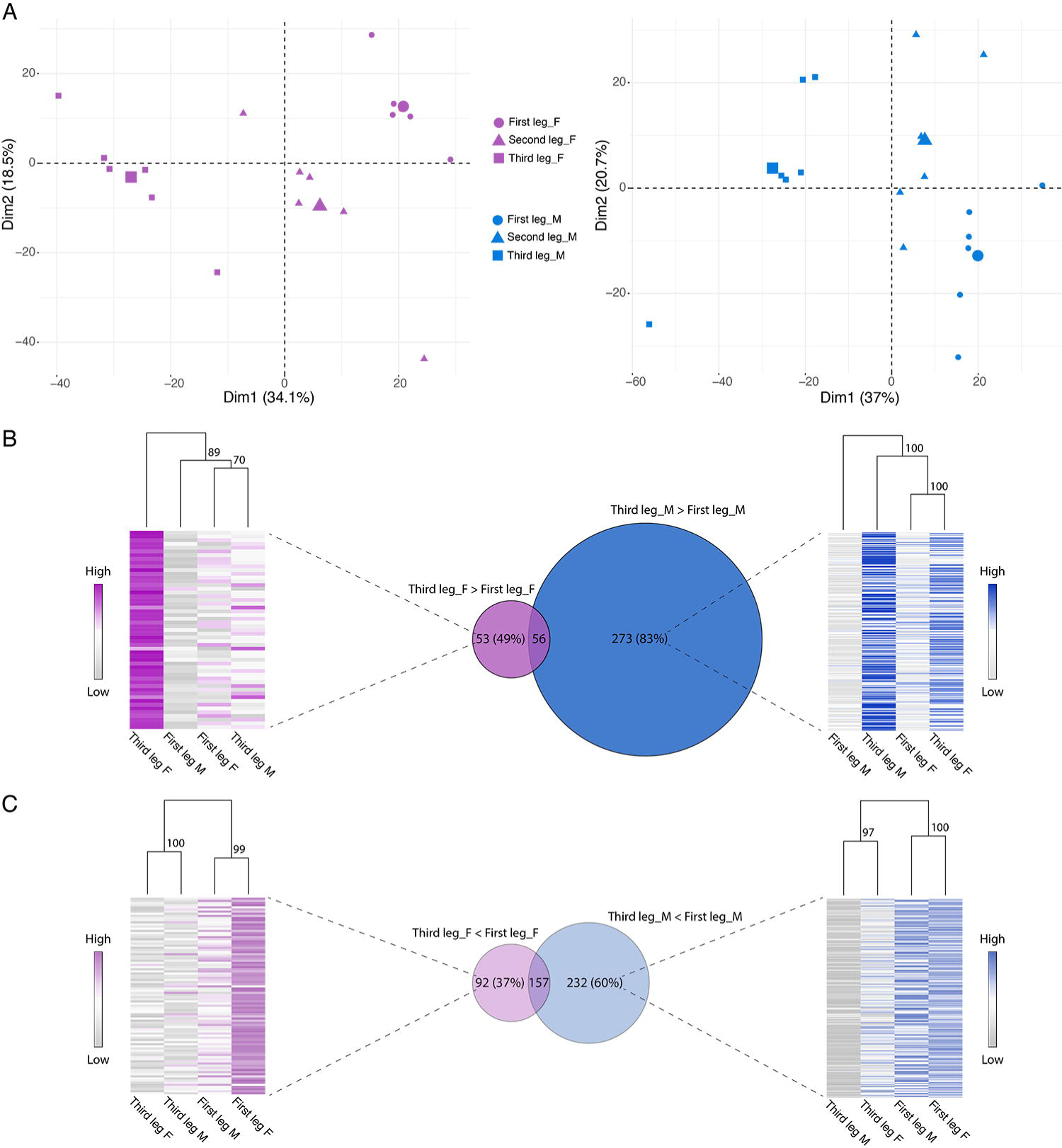
Signature of trait exaggeration among leg-biased genes. **(A)** PCA analysis of expressed genes in male and female legs separately. **(B-C)** Venn-diagrams illustrate the number of leg-biased genes shared in females (purple) and males (blue), both for up-regulated genes in the third **(B)** and first legs **(C)**. The size of the diagrams is proportional to the number of leg-biased genes (fold-change > 1.5). Heatmaps and hierarchical clustering show similarity in average expression between legs and sexes for the set of leg-biased genes specific to females (purple) and males (blue).

### Leg exaggeration is associated with a specific signature of leg-biased gene expression

Insect legs are serial homologs that share most of their developmental genetic program due to common origin ^23,24^. Therefore, we sequenced leg RNA, at the developmental stage where we detected a burst of growth, to match the observed morphological divergence in leg growth with differences in gene expression between serial homologs of each sex (Supplementary table 2). Although the differences in length between serial homologs were more pronounced in males than in females (Figure 1) ^21^, we failed to detect any correlation between such variation in morphology and variation in gene expression (Figure 2A). This is reflected in the first major axis of variation, which separated the legs with similar contribution (about 35% of the total variation) in both males and females (Figure 2A). This result suggests that morphological divergence in serial homologs between the sexes may not result from a global change in gene expression but rather from specific sets of genes.

To identify the genes underlying the growth differences in serial homologs, we examined the differentially expressed genes between the first and the third leg (leg-biased genes), and compared the divergence in expression between these two legs in males and females. We found that the third legs diverged more from the first legs in males than in females, both in terms of number of differentially expressed genes and in the degree of differential expression (Figure 2B; Supplementary figure 3). Similar results were observed for the second legs in males, which are moderately exaggerated (Supplementary figures 3 & 4). A hierarchical clustering analysis showed that the up-regulated genes restricted to male third legs (489 genes) are similarly expressed between the first and the third legs of females. Similar results were also observed for biased genes in female third legs (Figure 2B), indicating that a large proportion of up-regulated genes in the third legs are sex-specific. In the first legs, which have more similar scaling relationship between males and females, we found that about 63% of leg-biased genes were common between the sexes (Figure 2C). Moreover, hierarchical clustering analyses revealed that genes which were up-regulated in the first legs specifically in males showed similar expressions with the first legs of females (Figure 2C). Likewise, female-restricted genes showed similar expressions with the first legs of males (Figure 2C). Finally, leg-biased genes shared among sexes do not show any average differences in expression between males and females (Supplementary figure 5). Altogether, these data suggest that the morphological divergence of serial homologs is associated with up-regulation of a unique set of genes in the third legs of both sexes, with a higher magnitude, both in number and degree of differential expression, in male exaggerated third legs. We note also that the third legs express a considerable set of common genes between males and females, possibly underlying more global regulators of leg growth and allometry.

### Key developmental genes control the scaling relationship of male third legs

Extravagant signals and weapons, such as *M. longipes* third legs, are predicted to occur in good condition individuals because of the high fitness cost they impose ^7,25,26^. Proposed mechanisms for the development of exaggerated phenotypes include increased sensitivity or the production of growth factors by the exaggerated organ ^5,7^. To test this hypothesis, we examined the developmental genetic pathways that are enriched in the exaggerated legs through analyses of gene ontology in leg-biased genes (Supplementary table 3). Genes preferentially up-regulated in the third legs of males were enriched in metabolic processes, transmembrane transport and growth signalling pathways. Genes preferentially up-regulated in the third legs of females were enriched in signalling and metabolic pathways as well as response to stimuli and cell communication processes. Among the genes and pathways that were hypothesized to play a role in the regulation of growth-related exaggerated traits ^3^, many were identified in our datasets as differentially expressed between serial homologs within the same sex or between the sexes (Supplementary figure 6; Supplementary table 4; see Toubiana et al. companion paper for sex-biased genes).

To test the role of these genes in trait exaggeration, we conducted an RNA interference (RNAi) screen targeting about 30 candidates representing transcripts that were leg-biased, sex-biased, broadly expressed or tissue-specific (Supplementary table 5). This functional screen identified two genes, *Bone Morphogenetic Protein 11* (BMP11, also known as *Growth Differentiation Factor 11*) and *Ultrabithorax* (Ubx), with unambiguous and reproducible effect on leg exaggeration. BMP11 is a known secreted growth regulator with a systemic effect in vertebrates ^27,28^ (Supplementary Figure 7), whereas Ubx is a tissue-specific Hox protein known to be confined to the second and third thoracic segments in water striders ^29^. In our dataset, *BMP11* is expressed in all legs but was up-regulated in the third legs of males, whereas *Ubx* is absent from the first legs, lowly expressed in the second legs, and highly expressed in the third legs of males (Supplementary tables 2 and 5).

RNAi depletion of *BMP11* at the fourth nymphal instar significantly decreased the mean and variance of adult body and leg length (Figure 3A-B; Supplementary table 6 for statistical analyses). Importantly, RNAi against *BMP11* also significantly reduced the allometric coefficient of male adult legs (Figure 3C; Supplementary table 6). In comparison to the controls, *BMP11* knockdown individuals showed around 12% reduction in body length, and the allometric coefficient significantly dropped from 3.70 to 1.96 (anova/likelihood ratio tests, p-values < 0.05) (Figure 3B-C; Supplementary table 6; also see material and methods). The allometric coefficients in the first and second legs of males were also lower compared to their corresponding controls, but to a lower degree than in the third legs (Supplementary figure 8; Supplementary table 6). Therefore, *BMP11* increases both the allometric coefficient of all legs and mean body size in the males (Figure 3C; Supplementary figure 8; Supplementary table 6). Despite the high and differential expression of *BMP11* in the legs of females (Supplementary table 6), *BMP11* knockdown in females reduced body size but did not affect the scaling relationship between leg and body length (Figure 3A-C; Supplementary table 6). Interestingly, the effect of *BMP11* RNAi knockdown brings body size and scaling relationship of *M. longipes* males closer to its sister species *M. pulchella* (Figure 3A-C, Supplementary figure 8). Altogether, these results indicate that *BMP11* is a key growth regulator in *M. longipes*, playing a role in the regulation of both leg allometry and body size in males during development.

**Figure 3:**
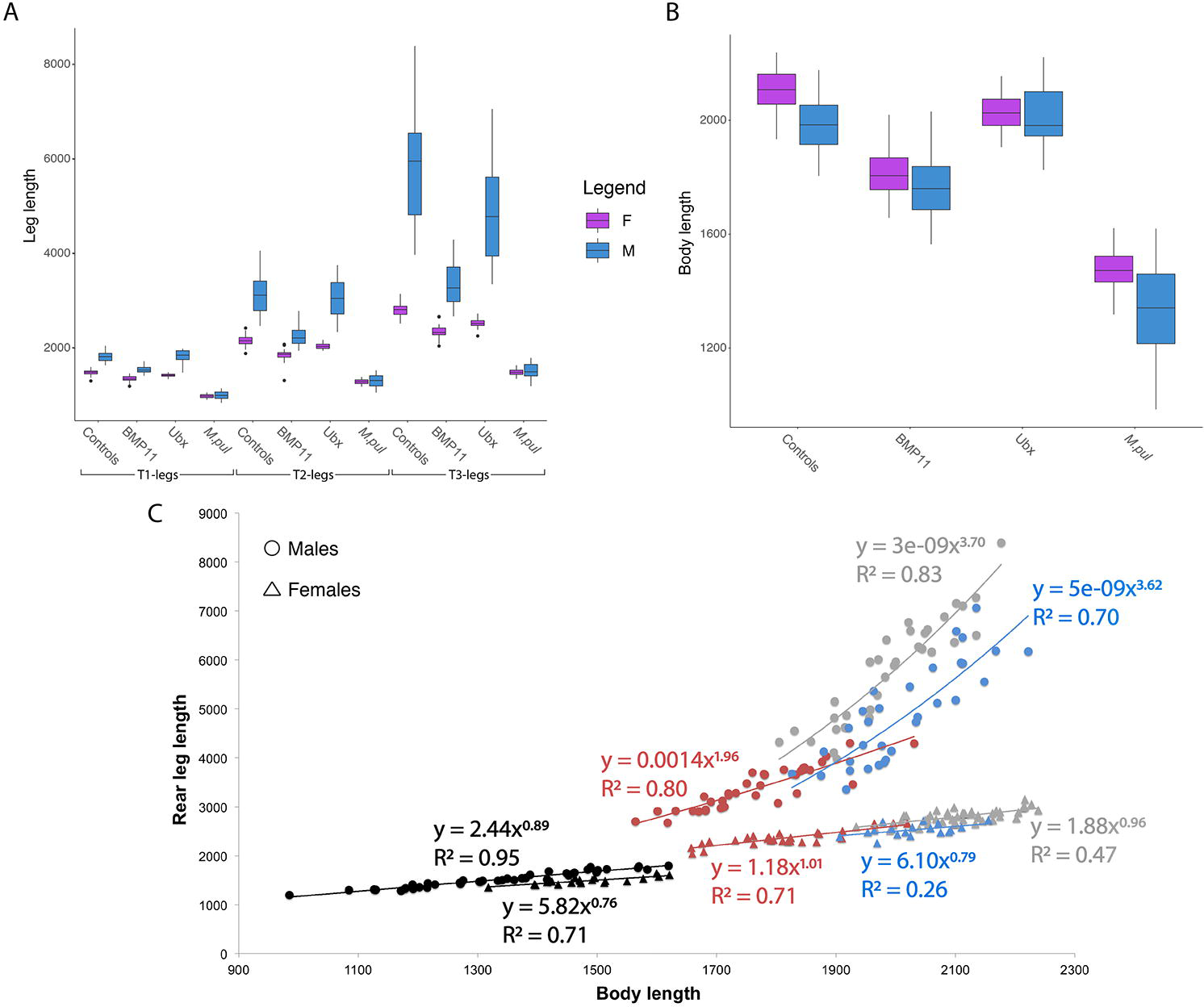
Effect of RNAi against BMP11 and Ubx on adult leg length (µm), body length (µm) and scaling relationships. **(A)** Effect of BMP11 and Ubx RNAi in adult male and female leg length (mean and variance) across the three legs. Statistical analyses can be found in Supplementary table 6. **(B)** Effect of BMP11 and Ubx RNAi on adult male and female body length. Statistical analyses can be found in Supplementary table 6. **(C)** Comparisons of adult static allometries between controls (grey), BMP11 RNAi (red) and Ubx RNAi (blue) in *M. longipes* males and females. Male and female static allometries of *M. pulchella* natural population (black) is also shown for evolutionary comparison. Power law regressions were fitted to the raw dataset. Statistical analyses were also performed on log-transformed data and can be found in Supplementary table 6.

Contrary to *BMP11*, *Ubx* knockdown did not affect the slope but rather shifted the intercept due to a 16% decrease in males’ third leg length (t-test: t = −3.5135, df = 62.904, p-value < 0.05) (Figure 3A-C; Supplementary table 6). We also detected a small effect in the second legs and no effect in the first legs, consistent with the expression of *Ubx* in these tissues (Figure 3A; Supplementary figure 8; Supplementary table 6). Contrary to *BMP11*, *Ubx* knockdown did not affect average body length (Figure 3B-C; Supplementary table 6). *Ubx* knockdown in females also altered the intercept in the third leg length allometry, but to a lower extent than in males despite the similar expression of *Ubx* (Figure 3C; Supplementary table 2 and 5; Supplementary table 6). This result first demonstrates that, in *M. longipes*, changes in leg length can be genetically decoupled from changes in body length. Second, this tissue-specific gene is associated with a change in intercept but not in allometric coefficient.

### *BMP11* knockdown males fight less frequently than control males

We have shown that the evolution of exaggerated third legs in *M. longipes* was accompanied with increased body size and increased male fighting intensity compared to its sister species *M. pulchella* (Figure 3; ^21^). To test whether *BMP11* also influences male fighting behaviour in *M. longipes*, we compared the frequency of fights between controls and individuals treated with *BMP11* RNAi in male-male competition assays (see Material and methods; Supplementary videos 1 and 2). Similar to the controls, *BMP11* RNAi-treated males attempted to dominate egg-laying sites, mated, and guarded females (Supplementary video1&2; ^21^). However, unlike the controls, these *BMP11* RNAi-treated males avoided fighting when other males approached egg-laying sites. Instead, they adopted a sneaking behaviour by focusing on the female rather than trying to chase rival males away, or by abandoning the site altogether without a fight (Supplementary video 2). We quantified this “docile” behaviour by calculating the frequency of fights on floaters and found that *BMP11* males fought on average twelve times less than control males (Figure 4A; t-test: t = 3.6134, df = 3.0559, p-value < 0.05). *BMP11* females were attracted by male’s signals on floaters, but we failed to observe any mating events during our trials (Supplementary video 2). We further observed that *Ubx* RNAi-treated males retained aggressive fighting behaviour (Figure 4A), although the fight between two rival males lasted significantly longer compared to controls (Figure 4B; Supplementary video 3). These results suggest that *BMP11*, but not *Ubx*, increases fighting frequency in *M. longipes* males. Collectively, our results show that in *M. longipes* males, *BMP11* regulates the development of the exaggerated weapon and the associated fighting behaviour.

**Figure 4:**
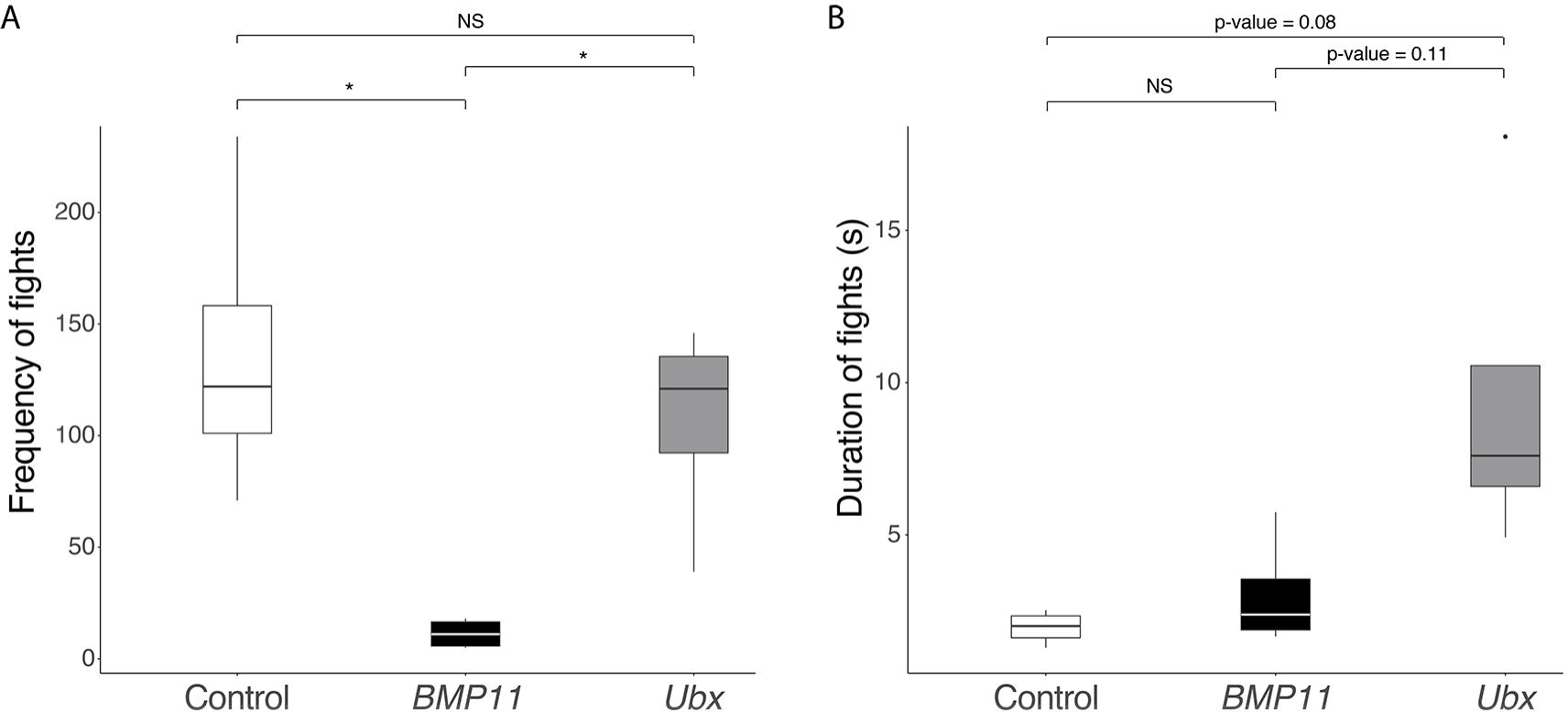
BMP11 and Ubx defects on male reproductive behaviour. **(A)** Defects of BMP11 and Ubx knockdowns on male fighting frequency compared to control individuals. **(B)** Defects of BMP11 and Ubx knockdowns on male fighting duration compared to control individuals.

## Discussion

We have shown that leg exaggeration in *M. longipes* males is associated with a specific signature of gene expression. Among the genes up-regulated in the exaggerated trait, we demonstrated that a broadly expressed growth factor, BMP11, increases slope, whereas a tissue specific factor, Ubx, increases intercept. Finally, we have shown that BMP11 regulates the growth of the body, the disproportionate growth of the legs, and also increases aggressiveness in males. This indicates that these three male traits are genetically correlated through the pleiotropic effect of BMP11. How BMP11 changes fighting behaviour in *M. longipes* males is unknown. A possible explanation is that the docile behaviour of BMP11-RNAi treated males can be due to a plastic response whereby males recognize their small body size and change reproductive strategy. Another possible explanation could involve a role of BMP11 in the development of nervous system components associated with the fighting behaviour.

Exaggerated traits represent striking cases of differential growth between various organs, and we are just beginning to understand the underlying developmental genetic processes ^5,30^. Studies in various insects suggest that traits would achieve exaggeration via their increased sensitivity to systemic growth factors such as Juvenile Hormone or Insulin ^7,10,11,31,32^. Our data also show that exaggeration, as seen in the third legs of *M. longipes* males, can be induced by growth regulating molecules other than hormones, such is the case for BMP11 and Ubx. These molecules are produced in excess by the exaggerated tissue, indicating an alternative path to trait exaggeration where over-production of the growth factor occurs locally rather than through heightened sensitivity to a systemic factor. Whether and how this increased tissue-specific expression of growth factors is connected to systemic growth pathways remains to be tested ^11^.

Our experiments describe a common effect of BMP11 on mean body length in both sexes. By contrast to males, the effect of BMP11 on body length is decoupled from its effect on leg length in females (Figure 5C). It is possible that increased body size has been favoured in both sexes, through increased competitiveness in males and increased fertility in females ^33,34^. Another explanation could be that smaller females were disfavoured due to reproductive incompatibility as body size increased in males. On the other hand, the lack of correlation in leg length between the sexes may be a consequence of sexual conflict resolution through dimorphism ^35^.

Studies of scaling relationships reported a strong stasis of the allometric coefficient at the microevolutionary time scale ^12–14^. Even at the macro-scale, slope is known to evolve slower compared to other scaling parameters such as intercept ^12,13^. Several hypotheses were formulated to explain such evolutionary stasis, including a lack of genetic variation, strong stabilizing selection acting on slope, or a constraining effect of pleiotropy ^12–15^. Our results, implicating BMP11 as a master regulator of growth in both body size and all three legs lend support to pleiotropy as a major driver of evolutionary stasis in allometric coefficient. In contrast, Ubx, which regulates the size of the third legs but not body size, has a role in changing the intercept. This suggests that the genetic architecture of slope may have more pleiotropic effect than that of intercept, possibly explaining the observed differences in evolutionary stasis between these two scaling parameters.

An important question is how trait exaggeration evolves despite the evolutionary stasis of slope. We have shown that the evolution of hyperallometry in *M. longipes* is associated with increased mean body size and male aggressiveness compared to its sister species *M. pulchella* ^21^. These changes are also associated with strong selection on males due to intense competition for access to females ^21^. In this context, selection is expected to favour traits that increase male competitiveness. Strikingly in *M. longipes*, BMP11 seems to regulate mean body size, allometric slope, and male aggressiveness, all of which increase male competitiveness. In this context, it is possible that directional selection favoured genotypes that regulate all three traits at the same time leading to the co-evolutionary process observed in *M. longipes*. Pleiotropy would therefore become both a promoting factor for the evolution of exaggerated weapons and a constraining factor involved in the stasis of slope.

## Material and methods

### Growth curves

*M. longipes* is a hemimetabolous insect that develops through successive moults which represent an exuvia of the previous nymphal instar. We raised each individual separately from 1^st^ nymphal instar to adulthood. Throughout this process, each individual produced five exuvia corresponding to the five nymphal instars. We then collected the exuvia and the adult from each individual and measured their leg lengths (Supplementary figure 1).

### Comparative transcriptomics: analyses of variance

Sample collection, transcriptome assembly and read mapping/normalization are described in a companion paper (Toubiana et al.). We performed an analysis of variance on sex separately (Figure 2). Here we took in consideration both line and replicate effects (See Toubiana et al. companion paper). The latter effect matched the days where RNA was extracted from each sample. We corrected for both effects, using Within Class Analysis, in subsequent analyses.

### Identification of leg-biased genes

The number of reads per “gene” was used to identify differences in expression among the different legs using DESeq2 ^36^. We first filtered transcripts for which expression was lower than 2 FPKM in more than half of the samples after combining the two inbred populations (12 samples total). Transcripts with average expression that was lower than 2 FPKM in the two comparing legs were also discarded. Finally, we ran the differential expression analysis by taking into account the line and replicate effects. We called differentially expressed genes any gene with a fold-change > 1.5 and a Padj < 0.05. All differential expression analyses were performed on the two lines combined as we aimed to identify genes involved in allometric slope, which is a common feature of both lines.

### Hierarchical clustering

Average expressions of leg-biased genes in the different sexes were clustered using Euclidean clustering in the R package PVCLUST version 1.3-2 ^37^ with 1000 bootstrap resampling. Heatmaps and clustering were performed using the log2(TPM) average expression of each gene from each tissue. Heatmaps were generated using the R package GPLOTS version 3.0.1.1.

### Nymphal RNAi

Double stranded RNA (dsRNA) was produced for *BMP11* and *Ultrabithorax*. T7 PCR fragments of the two genes were amplified from cDNA template using forward and reverse primers both containing the T7 RNA polymerase promoter. The amplified fragments were purified using the QIAquick PCR purification kit (Qiagen) and used as a template to synthesize the dsRNA as described in^38^. The synthetized dsRNA was then purified using an RNeasy purification kit (Qiagen) and eluted in Spradling injection buffer^39^ at a concentration of 6μg/μl. For primer information, see Supplementary table 5. Nymphal injections were performed in the line selected for long legged males^21^ at the 4^th^ instar as described in^40^. Injected nymphs were placed in water tanks, for which the water was changed every day, and fed daily with nine fresh crickets until they developed into adults. We measured adult legs, body, pronotum and mesonotum on a dissecting SteREO Discovery.V12 Zeiss stereomicroscope using the Zen Pro 2011 software. *BMP11* knock-down individuals had a significantly shorter pronotum compared to controls. This is evident in that the mesonotum is no longer covered by the pronotum (Supplementary figure 9). We used this effect on the pronotum to detect injected individuals that displayed wild type-like phenotypes, as is characteristic to the RNAi technique, by looking at the effect on the ratio pronotum-mesonotum (Supplementary figure 10). To further confirm that these individuals were unaffected by *BMP11* RNAi, we compared their scaling relationships to normal individuals and found that they display wild type-like phenotypes in all traits measured (Supplementary figure 11). We therefore removed those individuals from the final analyses.

### Behavioural assays

Male competition assays were performed using artificial puddles containing five egg-laying floaters. Each replicate corresponded to a population of five wild type females from the long-legged selected line^21^ and ten males from a treatment (either ten control or ten *BMP11*-RNAi males). Analyses were performed on four replicates for each condition. Male and female interactions were recorded on a Nikon digital camera D7200 with an AF-S micro Nikkor 105 mm lens. Video acquisitions were taken a couple of hours after the bugs were transferred to the puddle. We defined a fight as an interaction between two males going back-to-back and kicking with their third legs on or near a floater. If two contestants stopped fighting for more than five seconds, we counted the new interaction as a separate fight.

### Statistical analyses

All statistical analyses were performed in RStudio 0.99.486. Comparisons for the mean trait size and trait variance were performed on raw data, whereas log-transformed data were used for scaling relationship comparisons. We used standardized major axis (SMA) as well as linear models to assess differences in scaling relationships (‘smatr’ package and anova in R respectively, ^41^).

## Acknowledgements

We thank Francois Leulier, Severine Viala, Rajendhran Rajakumar for comments on the manuscript and Gael Yvert, Kevin Parsons and Roberto Arbore for discussions. This work was supported by and ERC Co-G WaterWalking #616346 and Labex CEBA grants for AK and a BMIC PhD fellowship to WT.

## Author contributions

AK and WT conceived the work, WT collected samples for sequencing, WT and DA performed analyses of gene expression, WT and AD performed RNAi screen, WT performed behavioural and statistical test, WT and AK wrote the manuscript.

