## Supplementary material for "Pleiotropy promotes male exaggerated weapon and its associated fighting behaviour in a water strider": Suplemental figures

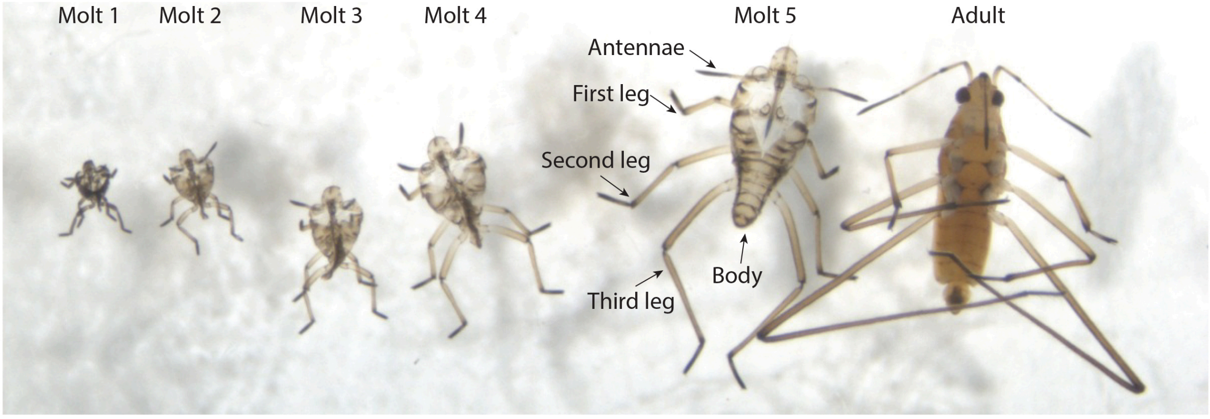


**Supplementary figure 1:** Representative picture of the nymphal molts left by an individual during its post-embryonic development in *M. longipes* (here a male). These molts were used to build a growth curve for each individual during nymphal development.

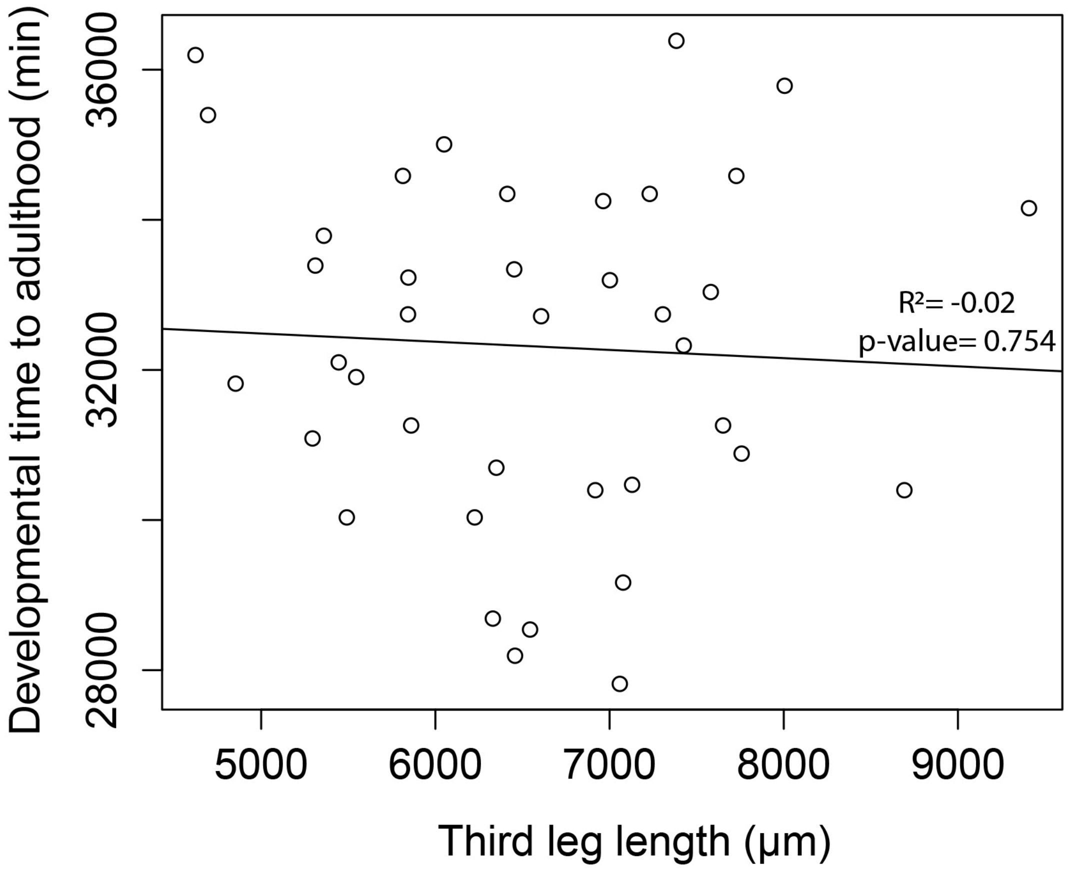


**Supplementary figure 2:** Covariation between male third leg length and duration of post-embryonic development (nymphal development) until adulthood. R-squared and p-value of the linear regression are indicated.

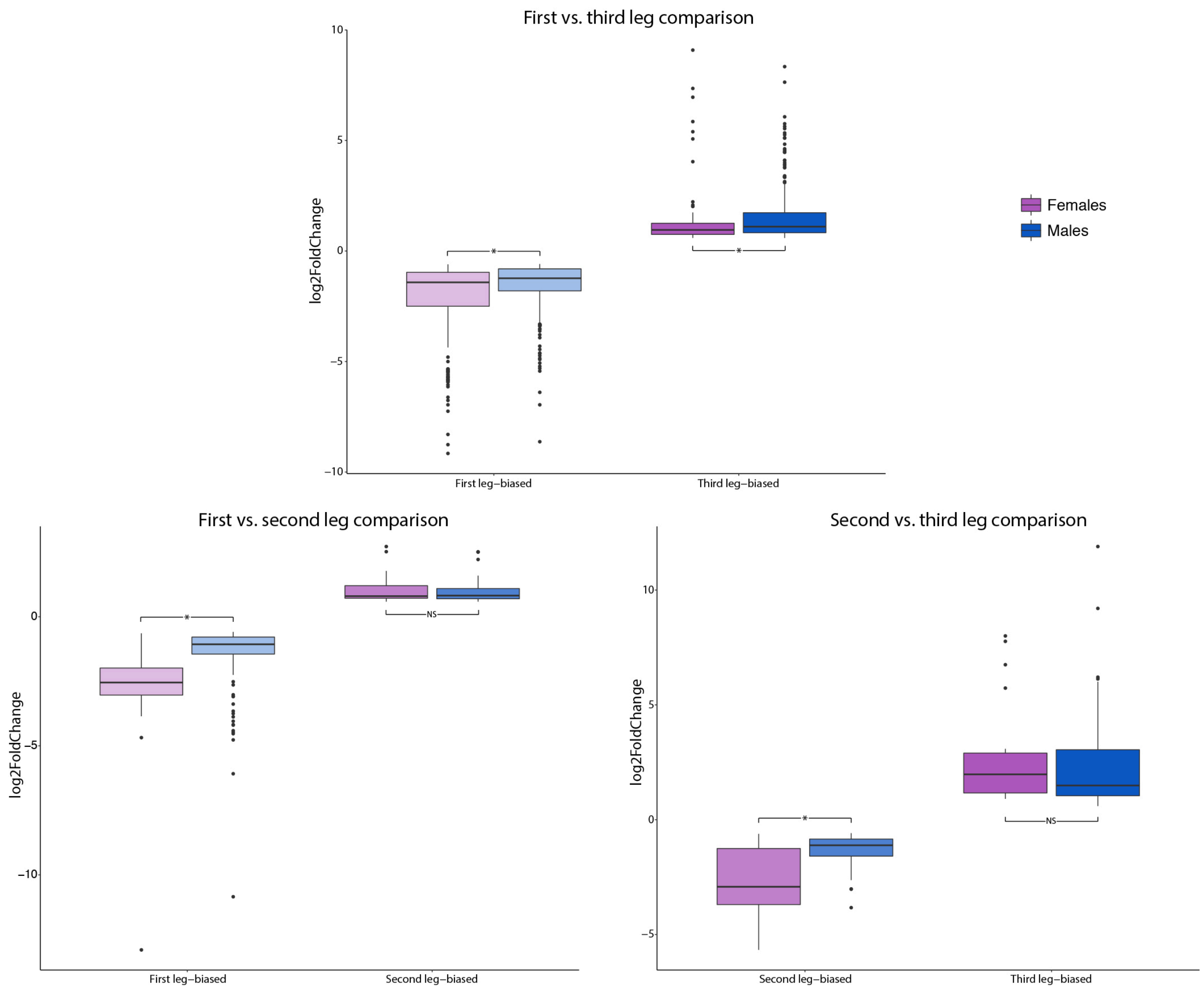


**Supplementary figure 3:** Boxplot differences in fold change (log2 Fold change) between males and females among the leg-biased genes.

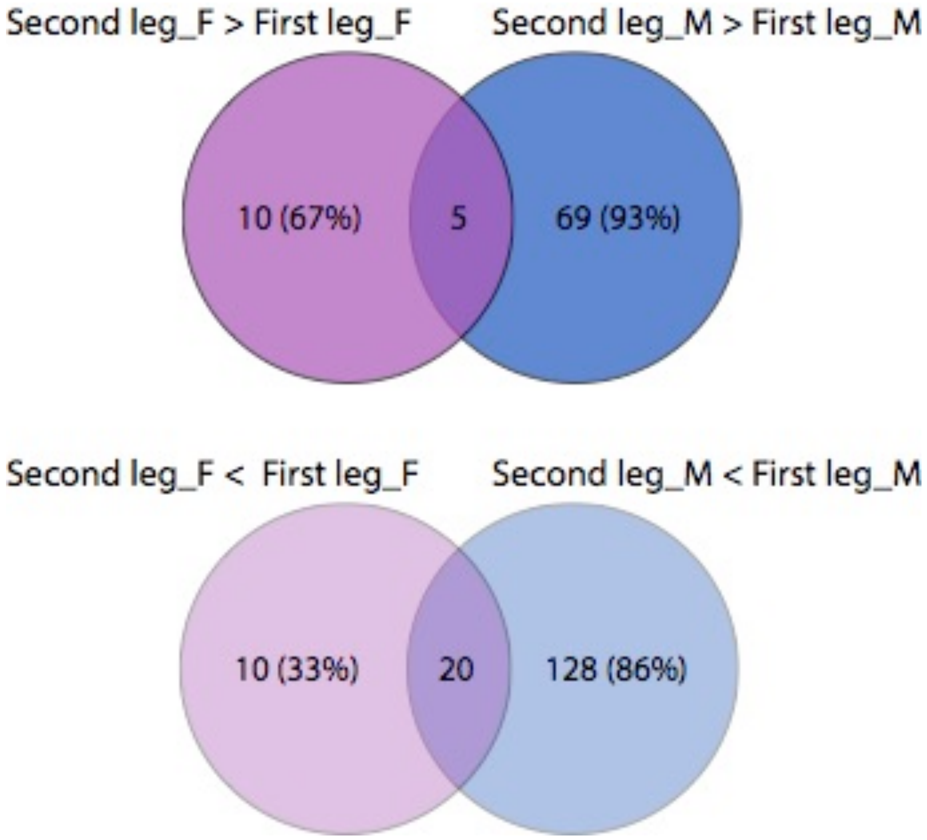


**Supplementary figure 4:** Venn-diagrams of leg-biased genes between the second and first legs between males (M, blue) and females (F, purple). Top are the upregulated genes in the second legs. Down are the upregulated genes in the first legs.

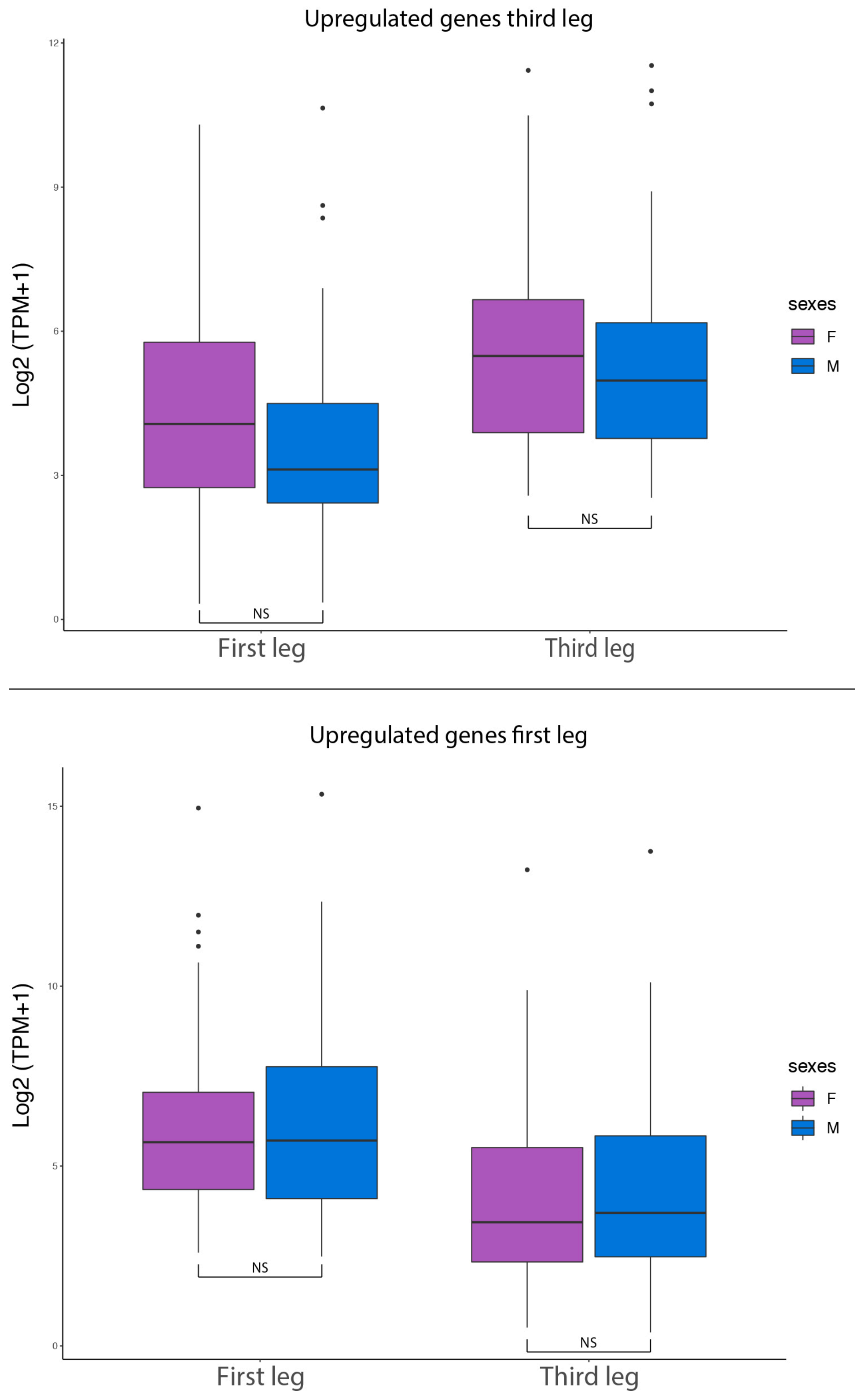


**Supplementary figure 5:** Boxplot differences in average expression (log2 TPM) of leg-biased genes in common between third and first legs of males and females.

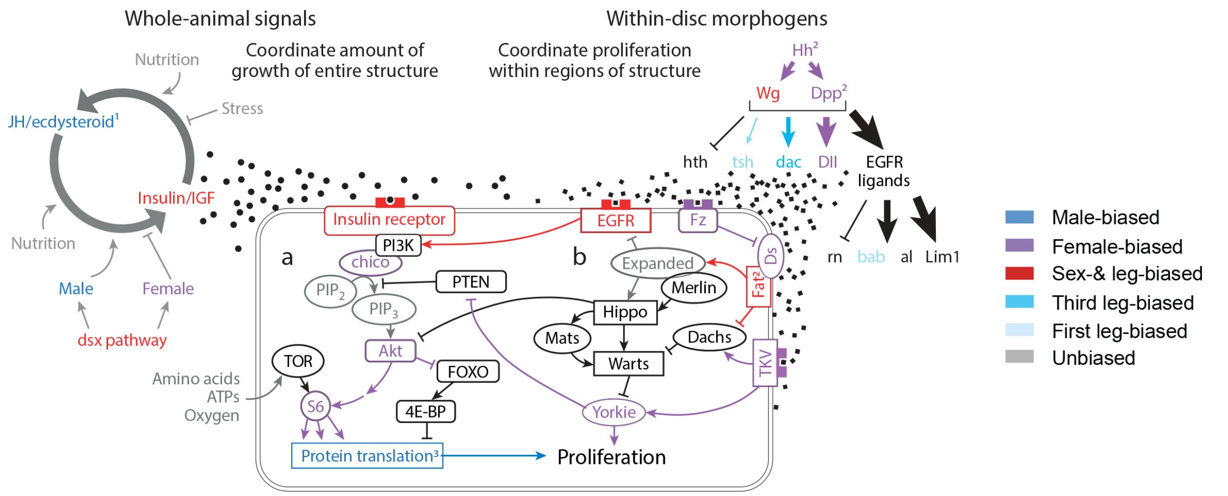


**Supplementary figure 6:** Pathways known or suspected to be involved in the regulation of exaggerated trait growth (from Lavine et al 2015). We color-coded the genes based on their expression pattern from the comparative transcriptomic analysis conducted in *M. longipes* (Toubiana et al. 2019). More details on each gene are found in Supplementary table 2.

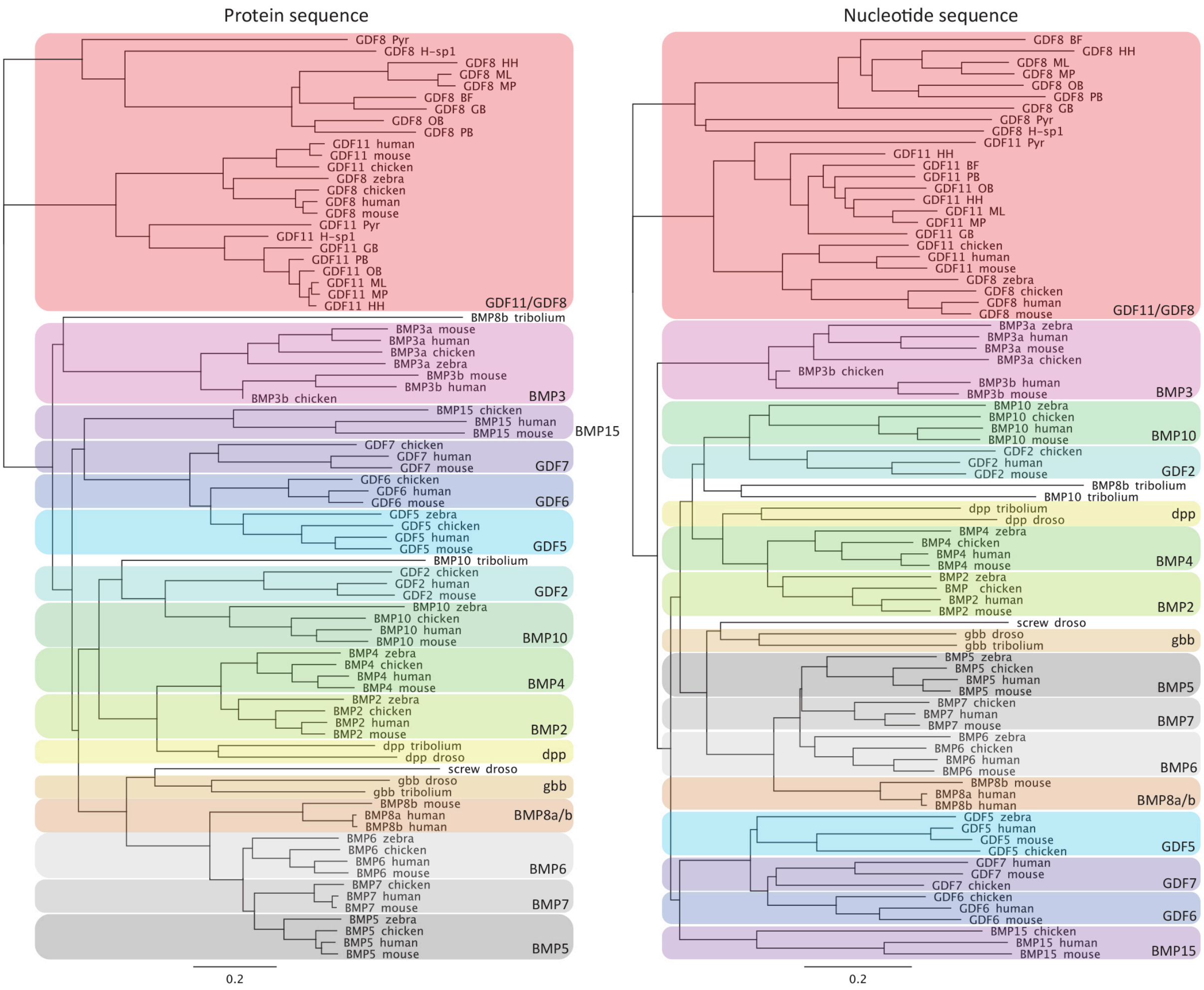


**Supplementary figure 7:** Phylogeny of BMP family (protein and nucleotide sequences) comparing BMP sequences in multipes species of water striders with vertebrates. BF: *Brachymetra furva*, HH: *Husseyella halophyla*, ML: *Microvelia longipes*, MP: *Microvelia pulchella*, OB: *Oiovelia brasiliensis*, PB: *Platyvelia brachialis*, GB: *Gerris buenoi*, Pyr: *Pyrrhocoris apterus*, H.sp1: *Hebrus sp1*, chicken: *Gallus gallus*, human: *Homo sapiens*, mouse: *Mus musculus*.

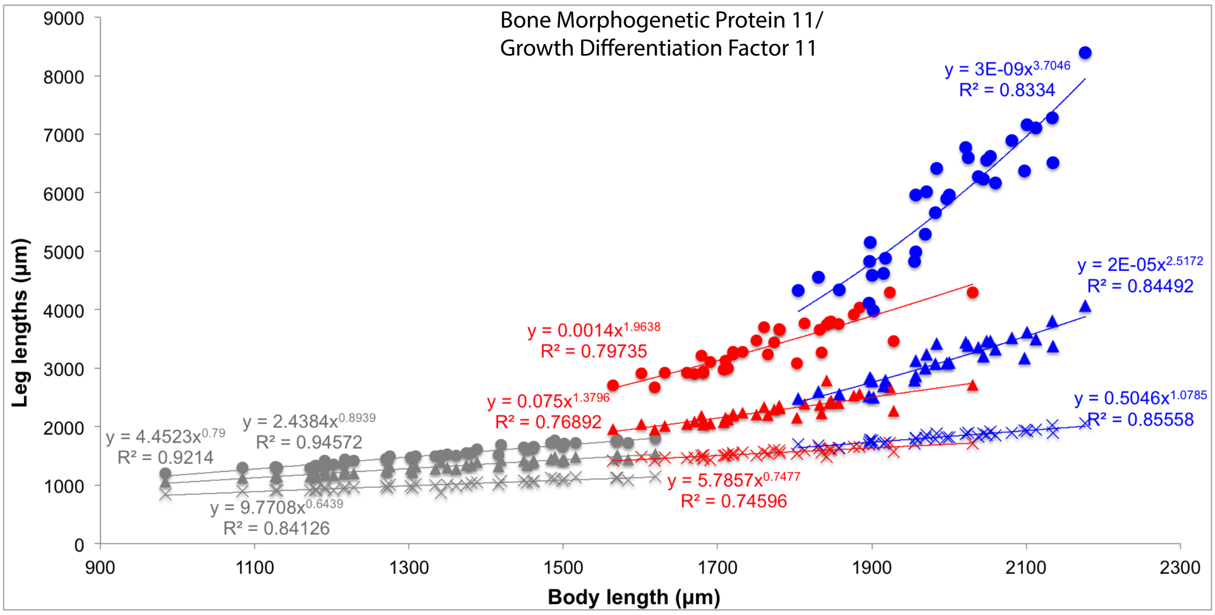
Bone Morphogenetic Protein 11/
Growth Differentiation Factor 11
Leg lengths (µm)
Body length (µm)
y = 3E-09x3.7046
R² = 0.8334
y = 2E-05x2.5172
R² = 0.84492
y = 0.5046x1.0785
R² = 0.85558
y = 0.0014x1.9638
R² = 0.79735
y = 0.075x1.3796
R² = 0.76892
y = 5.7857x0.7477
R² = 0.74596
y = 4.4523x0.79
R² = 0.9214
y = 2.4384x0.8939
R² = 0.94572
y = 9.7708x0.6439
R² = 0.84126

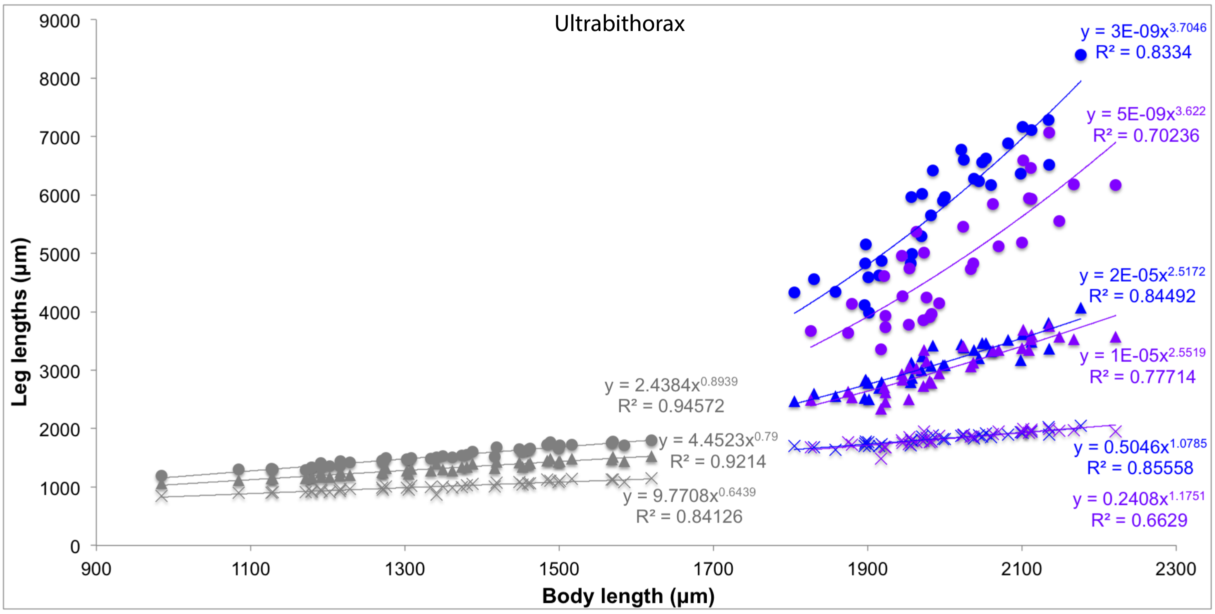
Ultrabithorax
Leg lengths (µm)
Body length (µm)
y = 3E-09x3.7046
R² = 0.8334
y = 5E-09x3.622
R² = 0.70236
y = 2E-05x2.5172
R² = 0.84492
y = 1E-05x2.5519
R² = 0.77714
y = 0.5046x1.0785
R² = 0.85558
y = 0.2408x1.1751
R² = 0.6629
y = 2.4384x0.8939
R² = 0.94572
y = 4.4523x0.79
R² = 0.9214
y = 9.7708x0.6439
R² = 0.84126

**Supplementary figure 8**: Static allometries of *BMP11* (red) and *Ubx* (purple) male knockdowns for all three legs. As comparison, we used control *M. longipes* individuals (blue) and *M. pulchella* natural population (grey). Circles correspond to third legs, triangles to second legs and crosses to first legs.

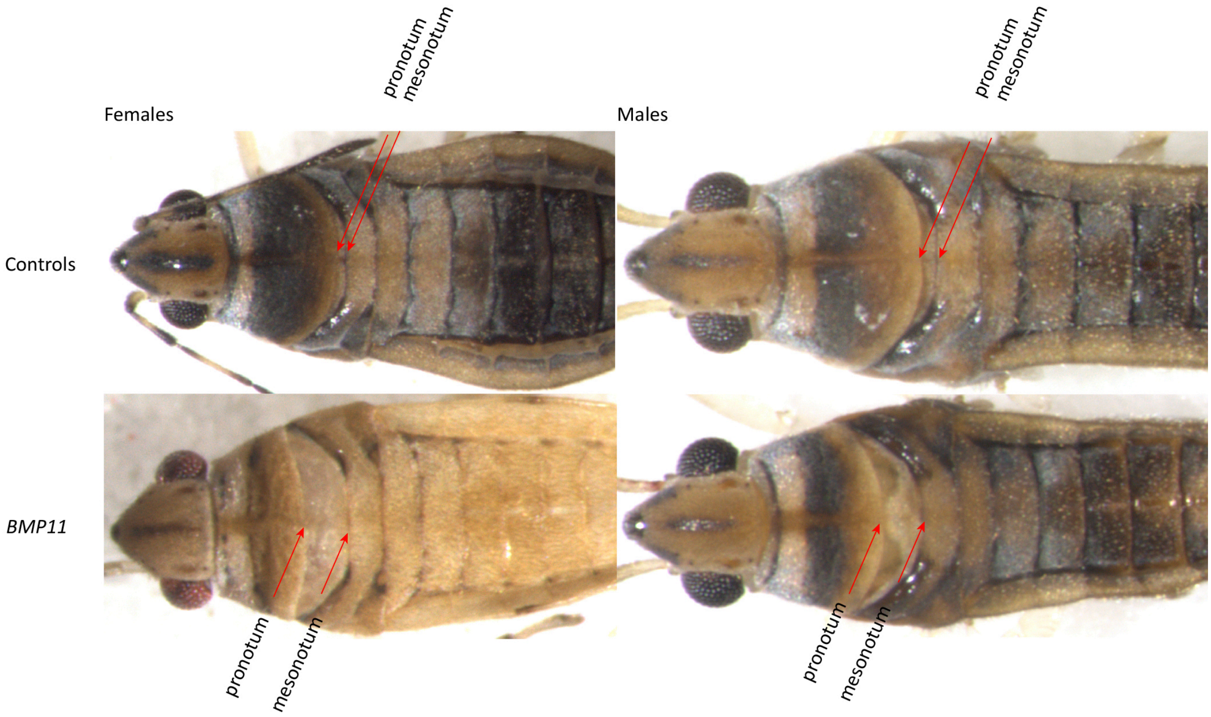


**Supplementary figure 9:** Effect of BMP11 RNAi on the growth of the pronotum indicated by red arrows. This effect was used to categorize individuals that were actually affected by BMP11 RNAi and those that were unaffected (see supplementary figure 9).

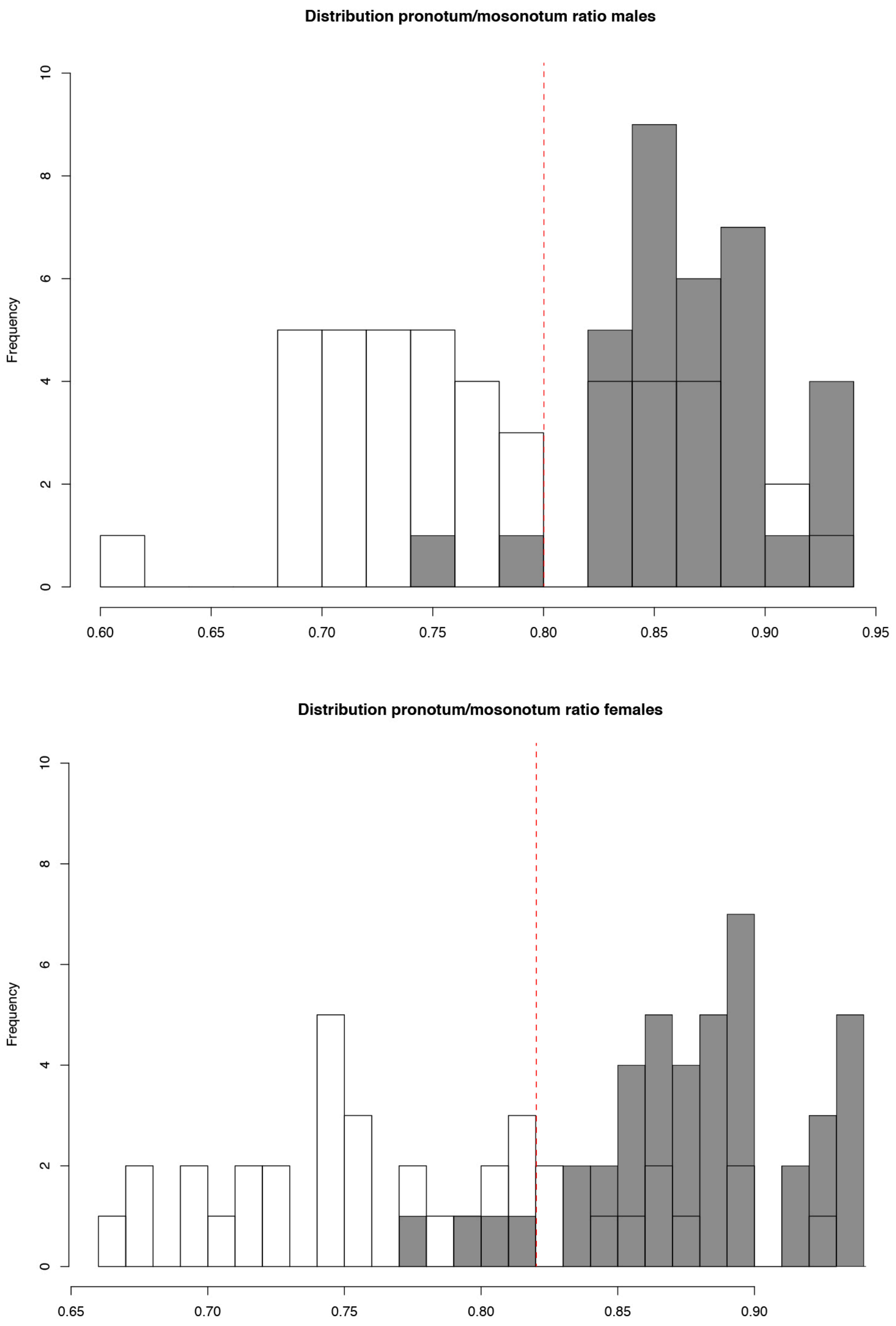


**Supplementary figure 10:** Distribution of pronotum/mesonotum ratios between controls and injected individuals with *BMP11* dsRNA in males and females. The red dotted line corresponds to the upper limit used to define an individual as *BMP11* knockdown. Other individuals were removed from the analysis, but see figure 10.

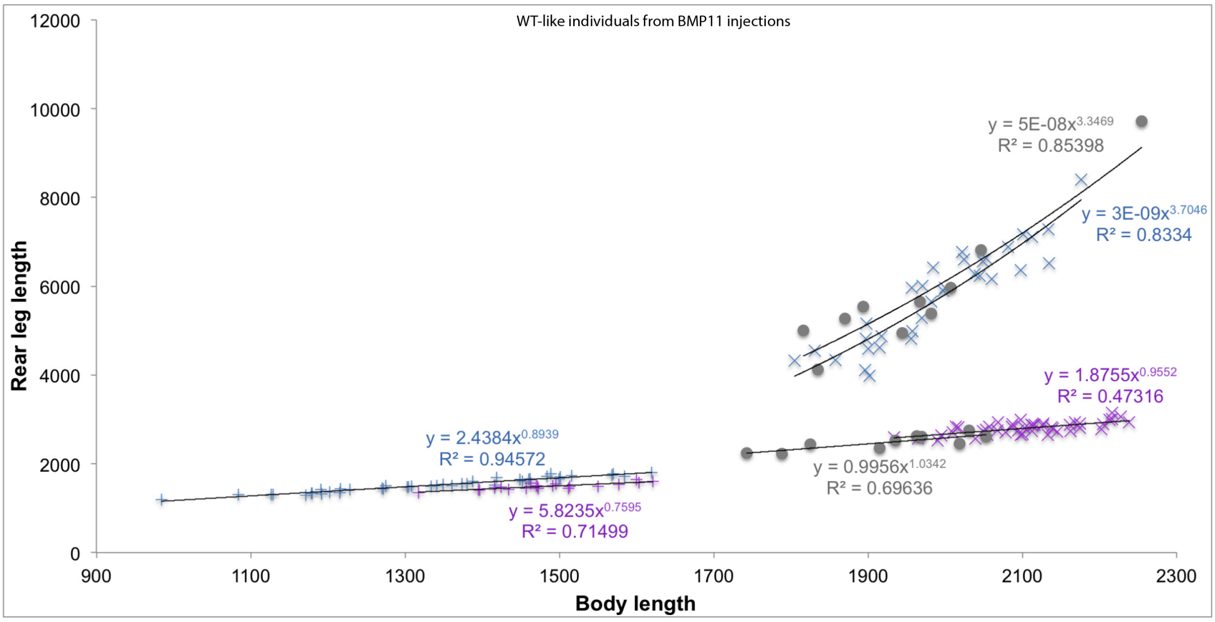


**Supplementary figure 11:** Static allometry of WT-like individuals after *BMP11* injection. The individuals categorized as WT (Grey spheres) based on the ration of pronotum/mesonotum (supplementary figure 9) show the same scaling relationships of leg to body length compared to controls individuals (x symbols).

**Supplementary table 1**: Ontogenetic leg allometry.

**Supplementary table 2**: Summary of raw read counts, TPM counts, blast analysis and the lists of leg biased genes across legs and sexes.

**Supplementary table 3**: Summary of Gene Ontology terms for leg-biased genes across legs and sexes.

**Supplementary table 4**: Summary of genes/pathways known or suspected to be involved in regulating exaggerated trait growth (from Lavine et al 2015).

**Supplementary table 5**: Identifiers, names, expression patterns, fold changes and phenotypes of the genes screened by RNAi.

**Supplementary table 6**: Summary statistics of *BMP11* and *Ubx* knockdown defects in leg lengths, body length and static allometries.

**Supplementary video 1**: Representative video of the fighting behaviour in *M. longipes* control individuals.

**Supplementary video 2**: Representative video of the fighting behaviour in *M. longipes BMP11* individuals.

**Supplementary video 3**: Representative video of the fighting behaviour in *M. longipes Ubx* individuals.
